# XL-MS–Guided Structure Prediction of Disordered *Encephalitozoon hellem* Proteins

**DOI:** 10.64898/2026.06.11.731626

**Authors:** Elizabeth Weyer, Yongliang Wang, Tadakimi Tomita, Carlos Madrid-Aliste, Prithviraj Nandigrami, Andras Fiser, Simone Sidoli, Jennifer T. Aguilan, Bing Han, Louis M. Weiss

## Abstract

Microsporidia such as *Encephalitozoon hellem* are obligate intracellular human parasites that remain genetically intractable, limiting functional characterization of their proteomes. Structural studies based on homology-based modeling and the use of deep learning algorithms of microsporidian proteins also remain limited because most have little to no sequence similarity to proteins with solved structures.

To address these limitations, we developed an approach that incorporates cross-linking mass spectrometry (XL-MS) data into structure prediction. XL-MS data provides upper bound distance constraints that can be incorporated into protein deep-learning based modeling and subsequent docking.

Using this approach, we generated a model for two interacting *E. hellem* spore wall proteins Spore Wall Protein 1B (Swp1b) and Endospore Protein 1 (EnP1), with no clear homologs outside of microsporidia, and which contain several disordered regions. These proteins are extremely abundant spore wall proteins of microsporidia and previously were not known to interact with one another. The resulting model not only is consistent with the experimental crosslinks used to generate the model but was subsequently confirmed by independently generated XL-MS data. The described AlphaLink-Modeller framework for structure prediction is particularly well suited to proteins with limited homology and/or substantial flexible regions, given they adopt a defined structural state within a biological context, thereby extending integrative modeling approaches to previously inaccessible targets.

## Introduction

Microsporidia are a large family of intracellular pathogens that evolutionarily are related to the Cryptomycota and are probably a sister taxon to the Fungi^1^. They are highly divergent and have undergone extensive genome reduction and rapid sequence evolution, resulting in proteins that are structurally and functionally distinct and frequently lack recognizable homologs in other organisms and for which few structures have been solved ^2,3^. Microsporidia are also genetically intractable, which limits the ability to use genetic modifications of genes to understand protein structure and function. In addition, many proteins in microsporidian invasion organelles of interest are comprised of a large percentage of disordered regions. As a result, the structure of many microsporidian proteins is difficult to predict, and it is also likely that X-ray crystallography would not be able to resolve the structure of these flexible regions.

Classical methods of protein modeling like homology modeling have struggled to predict structures of low-homology proteins, and *ab initio* modeling is limited by size. The AI boom has revolutionized protein modeling with the development of deep-learning based models such as AlphaFold^4^, RoseTTAFold^5^, DeepFold^6^, OmegaFold^7^, and ESMFold^8^. Millions of predicted protein structures are now available as models through AlphaFold DB^9^. However, it is notable that many of these predicted structures contain domains or whole proteins that are of low confidence. Out of the 49 model organisms that have their entire proteome predicted, approximately 30-40% of all residues have low or very low confidence scores (pLDDT < 0.7)^10^.This is, in part, because AlphaFold and other similar algorithms were trained on proteins with experimentally solved structures and large multiple sequence alignments^4^. Proteins with few to no homologs provide weak multiple sequence alignments, so predictions of these proteins are not as reliable^11–14^. Even single-sequence prediction algorithms like ESMFold that do not use multiple sequence alignments are less reliable for proteins with limited evolutionary signal and/or proteins with considerable flexible regions^11,15,16^. In general, flexible regions also do not usually yield defined structures when examined using currently established experimental methods such as X-ray crystallography and cryoEM^17,18^.

More recently efforts have been made to advance integrated structural modeling of proteins, where experimental data is combined with computational methods to determine the structures of proteins and their complexes^19–22^. Crosslink Mass Spectrometry (XL-MS) is a particularly useful tool to provide structural information for protein complexes with low homology and disordered regions. XL-MS uses MS-cleavable bifunctional cross-linking molecules that link two amino acids within a specific distance, thereby giving an experimental distance restraint on amino acid proximity^23^. These crosslinking molecules are added in the native protein environment, and so they capture these distance restraints in an unbiased fashion. Links formed between proteins (interlinks) provide an interactome between proteins in close proximity, and structural information on how two proteins interact. Links formed within individual proteins (intralinks) provide experimental distance restraints for the structures of the individual proteins in their native environment. These interlinks and intralinks are especially useful to inform modeling for proteins whose structures cannot be solved by traditional methods, and structures that struggle to be solved purely computationally. It has been integrated into existing tools like AlphaFold, docking pipelines, and atomistic simulations, making it generalizable for all proteins^24–31^.

Employing XL-MS to examine the proteome of *E. hellem*, we discovered that the proteins Spore Wall Protein 1B (Swp1b) and Endospore Protein 1 (EnP1) are in close proximity to one another as evidenced by the finding of numerous interlinks between these proteins in XL-MS experiments. Microsporidia have a thick spore wall that is made up of three layers – an inner plasma membrane, a middle layer of protein (endospore), and an outside layer of protein (exospore)^32^. EnP1 is a very abundant protein in the endospore. It is important to the structure of the wall but when secreted into the host cell it has been found to also have a role in modifying infected host cell gene expression^33,34^. Swp1b is a very abundant protein in the exospore and is involved in the adhesion of spores to host cells and in providing spore wall stability^35^. These proteins could interact with each other at the interface of the endospore and exospore layers in the spore wall.

Due to their high abundance and importance in the spore wall structure we were interested in modeling the interaction of Swp1b and EnP1, but these proteins are both low-homology proteins and are predicted to have a high percentage of flexible regions. Due to their predicted flexible regions and large size, the structure of these proteins was unlikely to be solved by x-ray crystallography, cryo-EM, or NMR. In fact, crystal production of heterologously expressed structural microsporidian proteins has, in general, not been successful. When Swp1b and EnP1 proteins are modeled using AlphaFold, the models are inconsistent with our experimental distance restraints (i.e., the cross-links identified by XL-MS both between these proteins and the internal cross-links within each protein). Even introducing distance restraints into an AI pipeline (AlphaLink) results in structures still incompatible with the experimental data obtained by XL-MS (i.e., all of the crosslinked peptides identified) Thus, we developed an approach that combines XL-MS guided AI modeling with XL-MS guided docking to provide structures that are consistent with the observed experimental distance constraints. In this paper, we present a novel method that allows one to model flexible, non-homologous protein complexes previously unable to be resolved with purely experimental or computational methods.

## Results

### Incorporating XL-MS data into modelling protein complexes

<FIGURE 1>

**Figure 1.**
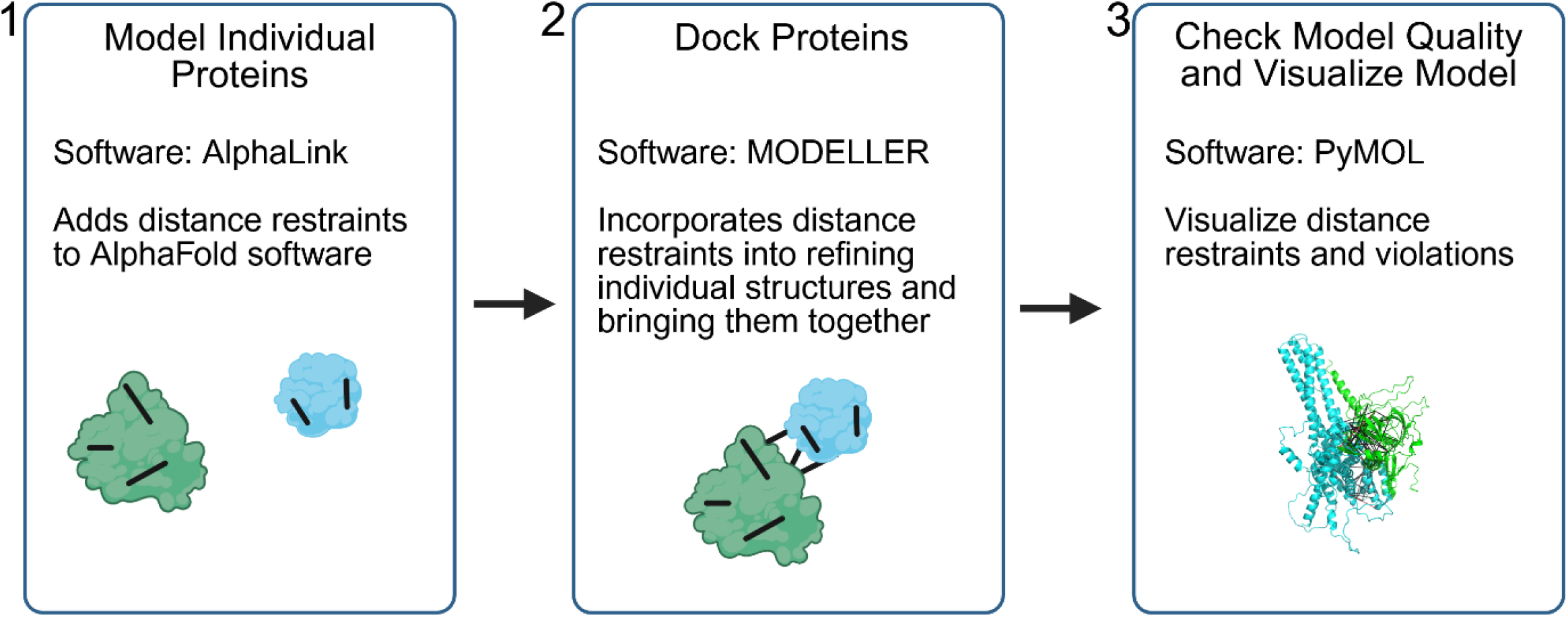
A new analytical approach for incorporating XL-MS data into modelling protein complexes. Each step of this three-step approach incorporates XL-MS data into its prediction, resulting in structures that are compatible with experimental XL-MS data, even proteins that have a large percentage of unfolded regions and no homology. Created with Biorender.com.

A new approach was designed to incorporate XL-MS restraints at each step of the modeling process. It is designed to work on proteins that have low homology to existing structures and have XL-MS restraints in their flexible regions. AlphaLink, a modified version of OpenFold that incorporates experimental restraints in structural prediction, is used to generate initial models of the monomers^26^.

The proteins are then brought together with MODELLER, a program that models target proteins based on templates by satisfying spatial restraints^36^. This step includes the intralinks present in the monomers to preserve their predicted structure while adjusting the structure to remove any distance violations, and the interlinks to bring the two proteins together.

This approach produces models that are consistent with experimental XL-MS data while preserving good structure quality metrics.

### Cross-linking mass spectrometry of *Encephalitozoon hellem* reveals proximity interaction of spore wall proteins Swp1b and EnP1

We were interested in using this approach to model proteins of *Encephalitozoon hellem*. Three independent replicates of crosslinking mass spectrometry experiments were performed on spores of *E. hellem* employing Azide-A-DSBSO, a mass spectrometer cleavable crosslinker spanning 14 angstroms that can crosslink nearby lysine, serine, threonine, and tyrosine residues. Previous studies indicate that the maximum expected cross linking distance cutoff for this crosslinker is 30Å, which incorporates the length of the crosslinker and amino acid side chains^37–39^. We identified 65 crosslinks with medium (FDR ≤ 0.05) and high (FDR ≤ 0.01) confidence between Swp1b and EnP1, demonstrating these proteins are in close proximity to one another. In addition, we identified 101 intralinks within EnP1 of medium and high confidence, and 58 intralinks within Swp1b of medium and high confidence (Figure 2a, Supplemental Data 1). Of these 224 crosslinks 97 were identical in at least two replicates (Supplemental Figure 1). To independently examine this protein-protein interaction, we performed co-localization analysis using anti-EnP1 and anti-Swp1b antibodies (Figure 2b). Fluorescence co-localization analysis revealed a stable interaction between Swp1b and EnP1 throughout spore maturation. Confocal imaging demonstrated complete overlap of Swp1b and EnP1 at all developmental stages, from early to mature spores. The yellow dashed lines in the merged images indicate the analysis paths. Line-scan profiles along these paths, obtained using ImageJ, showed perfectly synchronized fluorescence intensity peaks across all three channels, with Pearson correlation coefficients consistently above 0.90 (mean r = 0.94 ± 0.03). Negative controls using pre-immune mouse serum confirmed the absence of non-specific signals, further supporting the XL-MS data, which indicates that these proteins are in close proximity.

<FIGURE 2>

**Figure 2.**
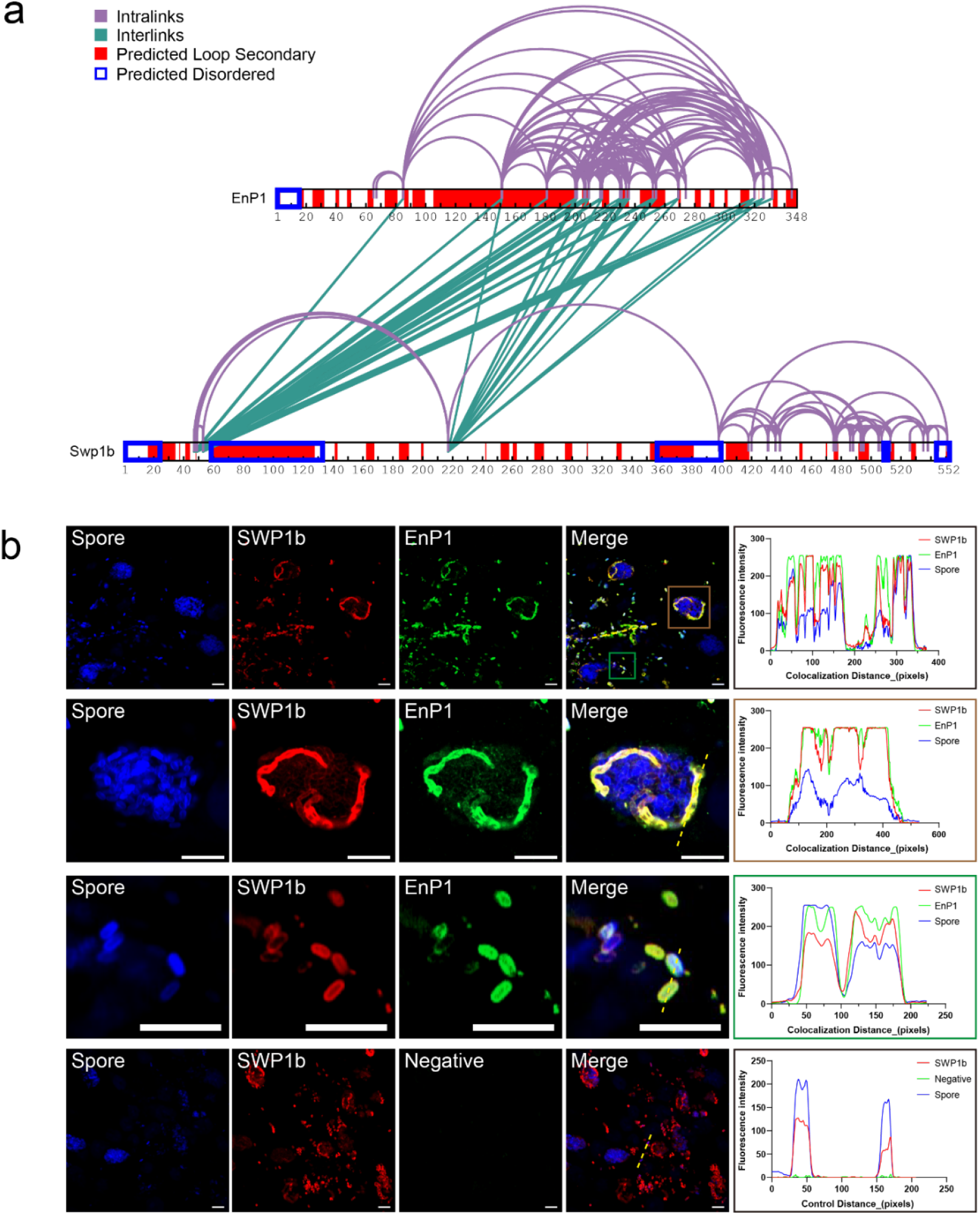
Swp1b and EnP1 are in close proximity to one another. **a**. Intralinks (purple) and Interlinks (green) mapped to amino acid sequences of Swp1b and EnP1. The location is indicated by the residue maps (Swp1b 552 amino acids, EnP1 348 amino acids). **b**. Swp1b and EnP1 interact during spore maturation. Confocal imaging shows that Swp1b (red), EnP1 (green) and spore (blue, fluorescent brightener) exhibit co-localization at different developmental stages. ImageJ line-scan profiles (right) reveal synchronized fluorescence intensity peaks, with Pearson correlation coefficients consistently above 0.90, indicating stable physical interaction between the two. Negative control: uninfected mouse serum. The yellow dashed line marks the analysis path in the merged image. Scale bar: 20 μm.

### Deep learning alone does not accurately model the Swp1b-EnP1 complex

Swp1b and EnP1 are both major components in the spore wall, making them a desirable target to model their structures and interaction with each other to better understand the whole spore wall architecture. However, this is made difficult by the fact that both Swp1b and EnP1 are highly divergent proteins with sequence homology largely restricted to microsporidia. BLASTp found homologs for both proteins exclusively in microsporidia, while PSI-BLAST identified only a single probable bacterial homolog for EnP1. Neither protein showed significant similarity to proteins of known structure using HHpred (Figure 3a). In addition, they both contain a significant portion of predicted flexible loop structure and disordered regions (Figure 2a). AlphaFold has become the most used algorithm for predicting unknown protein structures, however it must be used with caution. For example, in the case with Swp1b and Enp1, AlphaFold3 cannot predict the individual structures or the complex with confidence (pTM = 0.36, ipTM = 0.13). In addition to a low confidence pLDDT score in several areas of the protein (Figure 2b), the pTM and ipTM, metrics that measure the confidence in the overall predicted fold and position of the subunits indicate a likely failed prediction. Typically, when using AlphaFold3 we are reliant on the confidence scores alone to evaluate the accuracy of the model. XL-MS crosslink data, however, provides experimental distance restraints that can be used to evaluate the accuracy of the AlphaFold3 models. When we map the distance restraints (Figure 2a) to the AlphaFold3 structures, we see that this model fails to comply with 90/224 (40%) of the cross links for the complex, indicating that the AlphaFold3 predicted model is not compatible with the experimental structural data (Figure 3c). Even deep-learning algorithms like ESMFold that are designed to work on proteins with low homology do not produce structures compatible with our experimental distance restraints (Supplemental Figure 2).

<FIGURE 3>

**Figure 3.**
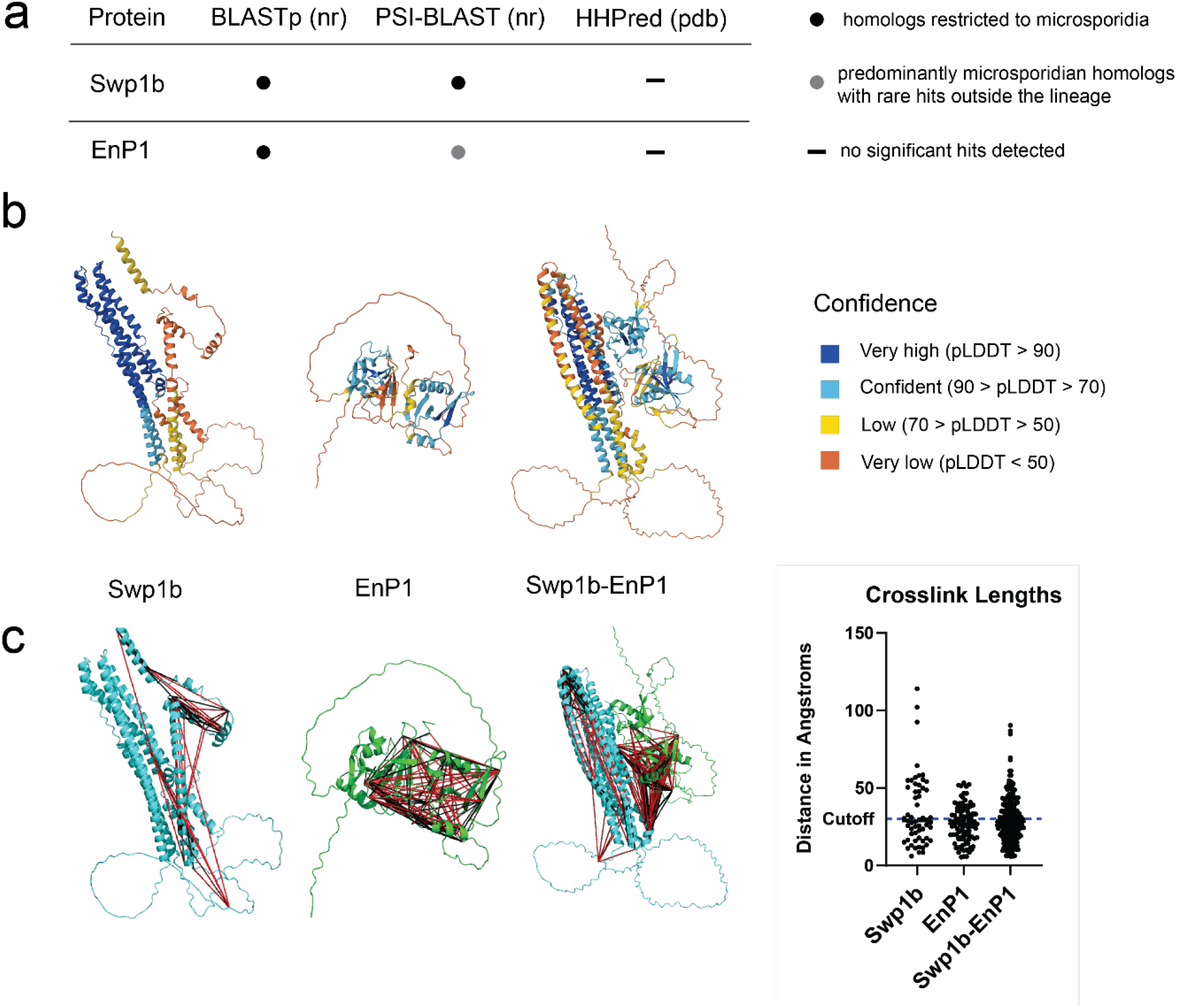
AlphaFold3 does not accurately predict the structure of Swp1b-EnP1. **a**. Sequence similarity searches were performed using BLASTp and PSI-BLAST against the nr database, and structural homology was assessed using HHpred against PDB-derived databases. **b**. Alphafold3 predicted models colored by pLDDT. **c**. Alphafold3 predicted models with crosslinks mapped onto the models with black lines. Distances are shown to the right, with the accepted cutoff of 30Å marked with the dashed blue line. Crosslinks with mapped distances above the 30Å cutoff are marked in red on the models.

### The AlphaLink-Modeller approach accurately models the Swp1b-EnP1 complex

AlphaLink, a modified version of OpenFold that incorporates experimental restraints in structural prediction, was used to generate initial models of Swp1b and EnP1. At this stage 77/101 (76%) EnP1 intralink distances and 41/58 (71%) Swp1b intralink distances are satisfied, compared to 61/101 (60%) EnP1 intralink distances and 32/58 (55%) Swp1b intralink distances satisfied in the AlphaFold3 model.

To enforce the full set of 224 intra and interlinks, the two chains were brought together with MODELLER by adding the crosslinks as additional distance restraints. This docked the two structures while preserving the global AlphaLink structures and removing any remaining distance violations.

When this approach is applied to the Swp1b-EnP1 complex, it produces a structure that is compatible with the experimental crosslinking data, with 208/224 crosslink restraints (93%) falling under the 30Å cutoff and the remaining 16/224 crosslinks falling under 31Å with an overall average length of 19.9 ± 6.5 Å. The structure contains most of its crosslinks in the interface between Swp1b and EnP1, indicating the greatest confidence in that area (Figure 4). The model predicts 10 hydrogen bonds and 7 salt bridges between Swp1b and EnP1 in the Swp1b-EnP1 interface (Supplemental Figure 3)^40^. The model was also assessed for overall global and local model quality. Specifically, we calculated RMS Z-score, which is a measure of assessing any inaccuracies in bond lengths and angles^41^. The results show reasonable quantitative measures with a bond length score of 1.166. bond angle score of 1.806, omega angle restraints of 1.487, and side chain planarity of 0.619, indicating no major deviations of bond and angular geometry from an optimized reference structure. Additionally, we assessed individual residue quality and composite model quality using QMEAN4 score, which also shows reasonable model quality at local residue level (Supplemental Figure 4)^42^. Direct comparison with known available structures is difficult because the model and complex we propose is unique, but these scoring metrics provide a reasonable model quality and validation.

<FIGURE 4>

**Figure 4.**
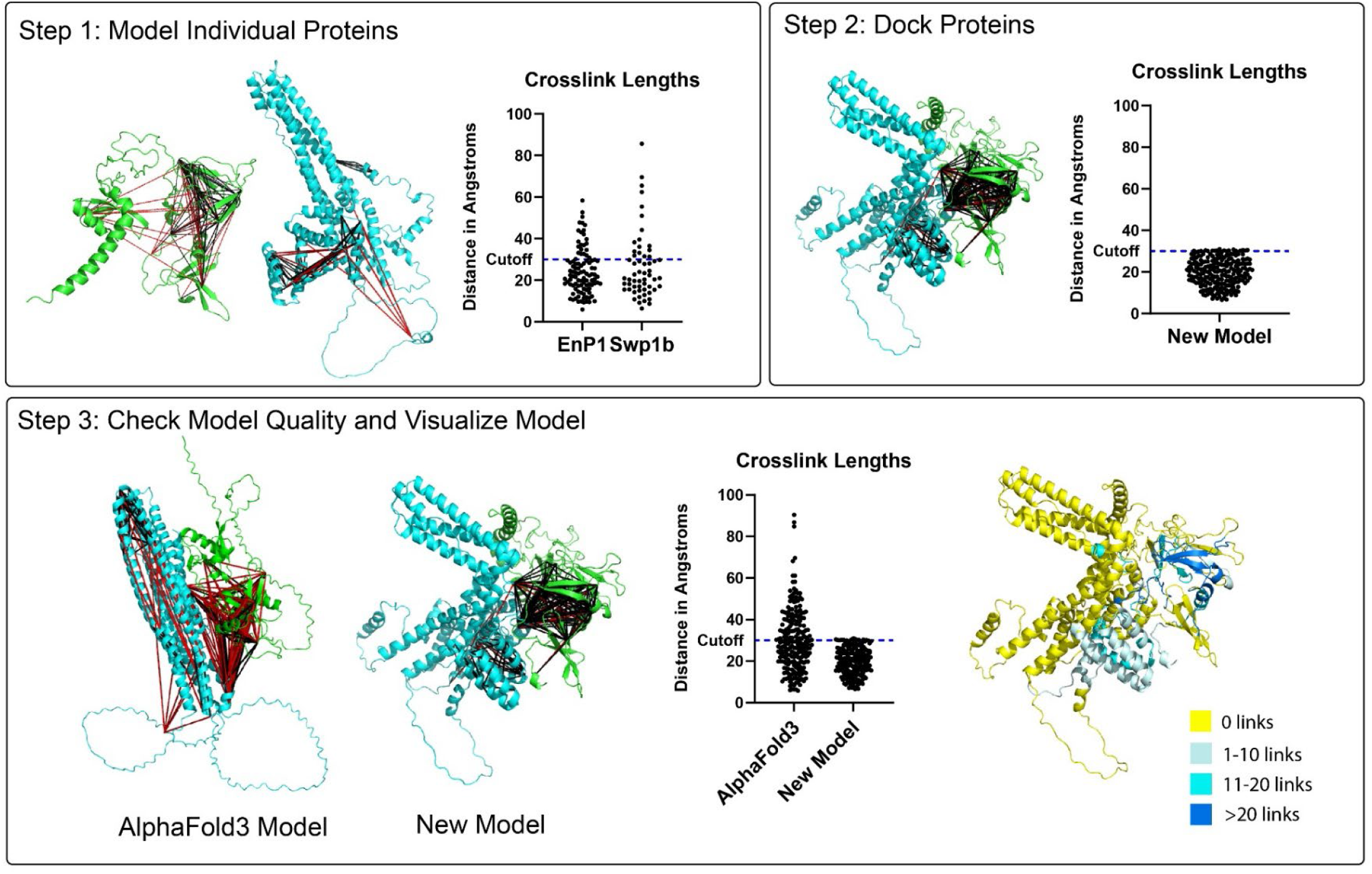
Stepwise modeling of Swp1b-EnP1 using the AlphaLink-Modeller approach. In Step 1 individual proteins are modeled with AlphaLink and crosslink distances are mapped on the right in black lines. Crosslink distances violating the 30Å cutoff are colored red. In Step 2 the individual proteins are brought together with MODELLER and the crosslink distances are mapped on the right. In Step 3 the model generated by AlphaFold3 (left) is compared to the model generated by our current methods (right). Crosslink lengths are mapped in the middle. To the right the structure is colored by the number of crosslinks in a 10 amino acid window, with yellow being 0 links and dark blue more than 20.

These models were generated with crosslinks combined from three separate XL-MS runs. To further test the resulting model, we performed a fourth small-scale XL-MS run and identified a total of 68 intralinks and interlinks in Swp1b-EnP1. This run was performed with DSSO, a different crosslinker that still has a distance cutoff of 30Å. 62/68 (91%) of these new links fell within the theoretical cutoff, with 4 additional links within only 1Å violation of the cutoff, 1 link at 39Å, and 1 link at 43Å. Of these DSSO links, 55 were identical to previously observed links and 13 were new links. Of the 55 identical links, 51 were within 30Å and 4 were within 31Å. Of the 13 novel links 11 were within 30Å, 1 was 39Å and 1 was 43Å (Figure 5, Supplemental Data 2). The link measuring 43Å is an EnP1 intralink. One end of the link is in an alpha helix with no other crosslinks having been previously identified in this area. All new links can be incorporated into the model without introducing any distance violations.

<FIGURE 5>

**Figure 5.**
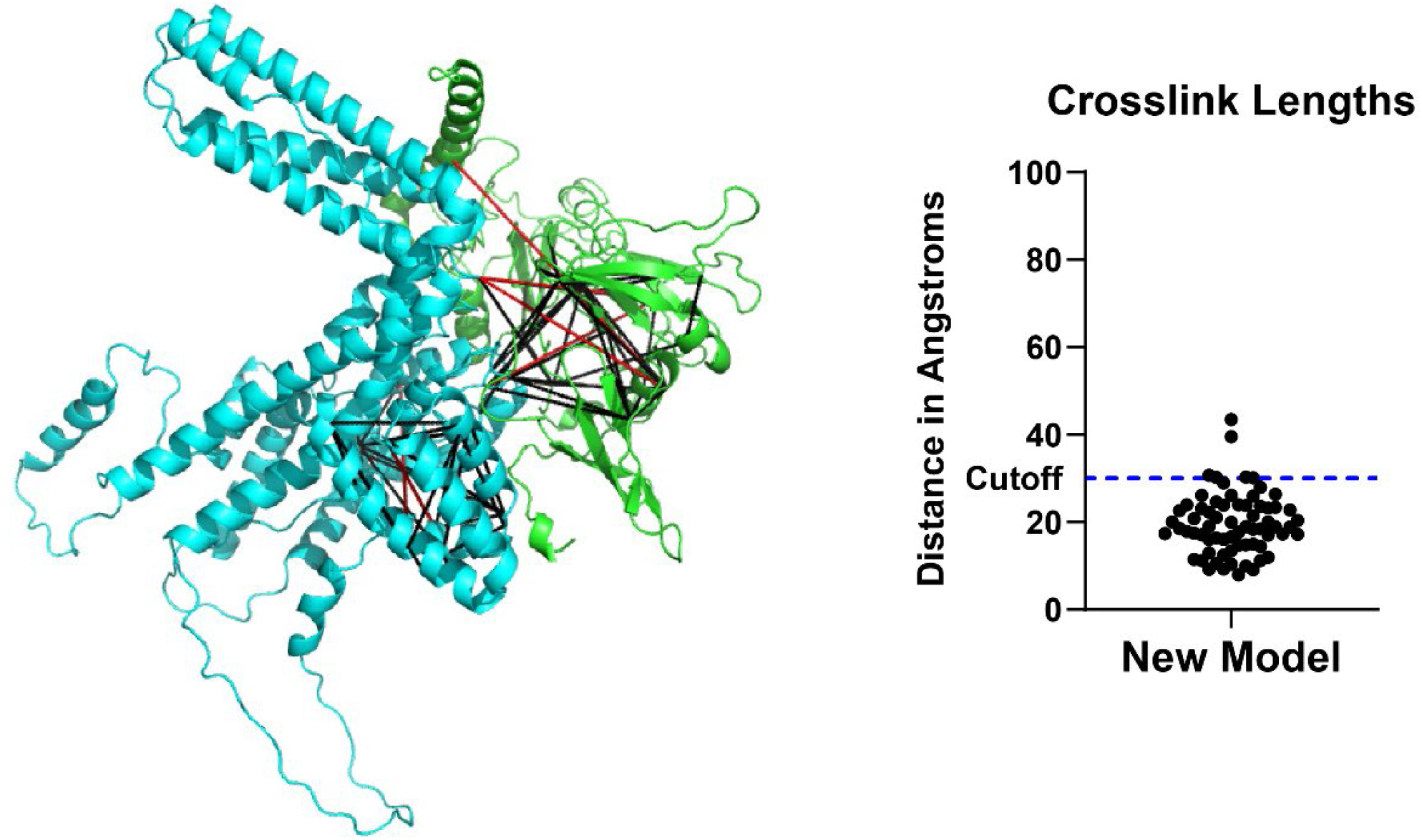
Independent data set maps onto the Swp1b-EnP1 model. Crosslinks generated from independent run mapped onto Swp1b-EnP1 model with links violating the cutoff colored red (left) and crosslink length distribution from independent run (right).

## Discussion

In this paper we use a new integrative modeling approach that incorporates XL-MS restraints into structure prediction of low-homology, flexible proteins. We used this pipeline to model a previously unidentified interaction between Swp1b and EnP1. While default deep-learning models failed to satisfy experimental XL-MS restraints, our integrative modeling approach produces models consistent with our experimental data.

Deep-learning protein prediction algorithms excel at predicting proteins with structural motifs already observed before^43^. But it struggles more when faced with proteins with unique sequence and flexible regions. Most deep-learning algorithms are trained to use multiple sequence alignments. Proteins without any detectable homologs lack this information and as a result structures predicted by Alphafold3 and similar algorithms are low confidence. Even deep-learning algorithms that do not rely on multiple sequence alignments still struggle with low homology proteins as they rely on sequence patterns as opposed to physical and chemical properties. As a result, predictions for low homology proteins are often of low confidence. In these cases, additional experimental data is needed to guide and evaluate modeling. Orthogonal experimental data such as XL-MS provides this type of experimental data to guide accurate structure prediction.

Modeling of Swp1b and EnP1 provides biological insight into microsporidia spore wall architecture. Prior transmission electron microscopy experiments localized Swp1b to the exospore and EnP1 to the endospore. Due to their location in separate layers of the spore wall, the modeling of this interface between the proteins reflects the structure of the interface between these two layers (Figure 6). It also demonstrates an example of flexible proteins becoming ordered during assembly. This data supports the broader concept that flexible regions can become structured in specific contexts, such as binding with a cognate partner. We believe this to be the case in Swp1b and EnP1 since individually they are predicted to have large regions of disordered regions and we were not able to detect this interaction using yeast-two-hybrid or co-IP experiments (Supplemental Figures 5 and 6) when these proteins were expressed in the cytoplasm of yeast, *Escherichia coli* or mammalian cells (HEK1). This argument is supported by cryo-EM images of the spore wall showing that it is highly organized, supporting regularly folded proteins instead of disordered proteins ^44^. This suggests that these proteins must be in the spore wall environment to fold correctly. This could be due to a number of factors, e.g. there are other spore wall proteins present in the exospore and endospore that may contribute to their folding correctly, and the endospore is largely made of chitin, which could act as a scaffold and influence EnP1’s structure in the spore wall. When modeling the two proteins together we can see they form a tight interface (Figure 6). We believe due to the abundance of these proteins and the location of their interaction that this complex is important for creating environmentally resistant spore walls. There are no genetic systems for Microsporidia (i.e. no transfection systems or methods to modify genes within these organisms have been developed), so we are not able to generate or test these models by altering genes within *E. hellem*. As this interaction is potentially important for the spore wall, our current model might serve as a starting point for drug development studies to disrupt this interaction.

<FIGURE 6>

**Figure 6.**
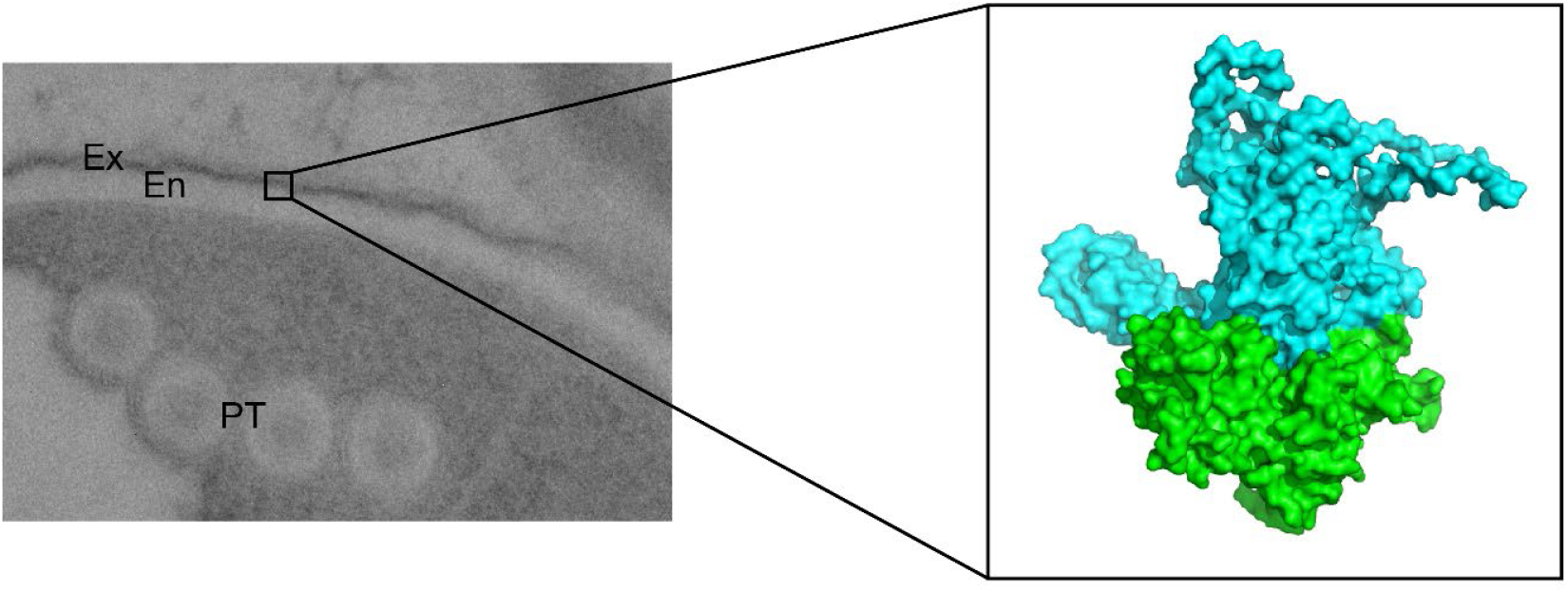
Model of the interface between the endospore and exospore. TEM image of the spore wall. Ex – exospore, En-endospore, PT-polar tube. The predicted model of Swp1b (blue) and EnP1 (green) form a tight interface we hypothesize is located at the interface between the exospore and endospore.

It is important to note the limitations of this approach. For example, this pipeline may not work for all disordered proteins. The approach is designed to identify a mostly ordered state, whether that is due to environmental context or binding with a partner.

This approach also does not intrinsically know the stoichiometry of an assembly. AI models tend to be able to better predict assemblies when the stoichiometry is known, which must be clarified with other experimental methods.

The results presented illuminate the value of integrative approaches that combine AI-based and physics-based predictions with experimental data such as XL-MS for modeling proteins that are experimentally inaccessible and can be precited with low confidence only using default approaches. As structural biology moves toward defining protein machinery by the incorporation of ‘omics’ level data, the use of this type of analytical pathway that can use such data to refine structural predictions is obvious and will be an essential part of computational modeling that aims to derive biologically meaningful structures.

## Methods

### Cell Culture

*E. hellem* were grown in Human Foreskin Fibroblast cells (HFFs; ATCC: CRL-1634; Hs27) in Dulbecco’s Modified Eagle medium supplemented with fetal calf serum, L-glutamine, and penicillin-streptomycin at 37ºC 5% CO_2_. Spores were passaged to new HFFs once a week. The spores in the supernatant were then collected after lysing out of the HFFs. After spinning down at 2000 x g for 20 minutes, they were then resuspended in 30 mL PBS and put through at 27-gauge needle thrice and filtered through a 5-uM filter (Millipore) once. They were then centrifuged at 2000 x g for 20 minutes and resuspended in 15 mL 1% H_2_O_2_ in PBS. They were then incubated at 37ºC rotating overnight to allow them to germinate. The spores were then centrifuged at 2000 x g for 20 min and washed three times with distilled water.

### Cross-linking Mass Spectrometry of *Encephalitozoon hellem*

The crosslinking protocol was performed following our previously published method ^45^. However, as the crosslinking protocol for *E. hellem* was being optimized at the time, all three runs had slightly different parameters. Changes were as follows. In Run 1, 5 150 mM plates of HFFs were used. The crosslinker was incubated for 4 hours at room temperature. Proteins were reduced with 130 mM TCEP, alkylated with 260 mM IAA, and digested with 1:25 trypsin over 24 hours. 20 µL of DBCO beads were added per mg of initial protein. No high pH fractionation was performed. In Run 2 15 150mM plates of HFFs were used. The crosslinker was incubated for 1 hour at room temperature. Proteins were reduced with 130 mM DTT in 5% sodium deoxycholate, alkylated with 260 mM IAA, digested with 1:25 LysC 37C overnight, then digested with 1:10 trypsin at 47ºC for 1 hour. 20 µL DBCO beads per 1 mg measured peptide concentration were added. No high pH was performed. Run 3 had the same parameters as run 2 except high pH fractionation was performed as previously described^45^.

Run 4 was performed as a test of a new method. Germinated spores were crosslinked with 1 mM DSSO for 30 minutes shaking room temperature. Proteins were reduced with 130 mM TCEP at 70ºC for one hour and alkylated with 260 mM IAA for 30 minutes room temperature. Phosphoric acid to 1.2% v/v was added to the solution, and the sample was loaded on a midi S-trap binding column. It was washed with S-trap binding buffer, and LysC was added 1:25 and digested for four hours at 37ºC. 1:25 trypsin was added and digested overnight at 37ºC. Peptides were eluted as per S-trap instructions, with an extra elution step of 90% acetonitrile 0.1% TFA, and peptides were dried in a speed-vac. The peptides were then resuspended to 10 mg/mL in 30% acetonitrile 0.1% TFA. 1.7 mg of peptide were injected into a Superdex 30 Increase 10/300 column (Cytiva), run with 0.5 mL/min flow rate, and eluted isostatically with the same buffer. 100 µL fractions were collected. Fractions from the first six peaks were selected and dried.

### Mass Spectrometric Analysis

Mass spectrometric analysis for all runs was performed as previously described ^45^. Samples were resuspended in 0.1% TFA and loaded onto a Dionex RSLC Ultimate 3000 (Thermo Scientific), coupled online with an Orbitrap Exploris 480 (Thermo Scientific) with FAIMS Pro attached. Chromatographic separation was performed with a picofrit analytical column (75 µm ID, 25 cm length) packed in-house with reversed-phase Repro-Sil Pur C18-AQ 3 µm resin. Peptides were separated using a 180-min gradient from 4% to 30% Buffer B (buffer A: 0.1% formic acid, buffer B: 80% acetonitrile + 0.1% formic acid) at a flow rate of 300 nL/min. The mass spectrometer was set to acquire spectra in a data-dependent acquisition mode. Briefly, the full MS scan was set to 375-1600 m/z with a resolution of 120,000 (at 100 m/z). MS/MS was performed at 60,000 resolution and AGC target of 1 × 10e4 and stepped HCD normalized collision energy of 19%, 25%, and 32% with FAIMS compensation voltages of −50, −60, and −75^46–49^.

The data were analyzed with Proteome Discoverer 2.5 with IMP-MS2 spectrum Processor for deisotoping and MS Amanda 2.0 with the following parameters: maximum missed cleavage 4, MS1 tolerance 10 ppm, MS2 tolerance 20 ppm, static modification: carbamidomethyl 57.021 Da (C), and dynamic modifications: oxidation 15.995 Da (M), DSBSO Amidated 325.065 Da (K), DSBSO Hydrolyzed 326.049 Da (K),. Spectrum searches were performed on MicroporidiaDB (microsporidiadb.org) release 65^50^. Data sets were analyzed with MS Annika 2.0 as MS-cleavable crosslinkers with crosslink doublet 308.039 on K, S, T, and Y residues^51^.

The fourth data set was analyzed with MS Amanda 2.0 with parameters DSSO Amidated 175.030 Da (K) and DSSO Hydrolyzed 176.014 Da (K). It was analyzed with MS Annika 2.0 as MS-cleavable crosslinkers with crosslink doublet 158.004 on K, S, T, and Y residues.

### Data Processing

The data generated by Annika 2.0 were processed using R with tidyverse packages. FDR was recalculated using a custom R script^45^. Peptide pairs are collapsed to residue-to-residue pairs. Quality scores are adjusted from lower-level scored by calculating Euclidean norms, and the FDRs were recalculated at higher levels after inter/intra crosslinks were separated according to the previously described method ^52^. Crosslinks were combined from all three runs. Crosslinks appearing in multiple runs were weighted accordingly (crosslinks found in two runs occurred two times, crosslinks found in three runs occurred three times). When analyzing distances only unique crosslinks were calculated. Links between Swp1b and EnP1 were visualized with xiVIEW (https://xiview.org/index.php)^53^. Secondary structure prediction was performed with PSIPRED (https://bioinf.cs.ucl.ac.uk/psipred/).

### Immunofluorescence staining of Swp1b and EnP1

HFF (ATCC CRL-2522) cells and *E. hellem* were maintained in DMEM (Gibco) supplemented with 10% fetal bovine serum (FBS) and 1% penicillin-streptomycin (Gibco) at 37°C under 5% CO_2_. For experiments, cells were seeded at 5 ×10^4^ cells/well on glass coverslips in 24-well plates. Fixed with 4% paraformaldehyde (PFA) for 15 min at room temperature (RT), cells were permeabilized with 0.1% Triton X-100 (15 min, RT) and blocked with 5% bovine serum albumin (BSA; Sigma-Aldrich, A7906; 1 h, RT). Primary antibodies were applied overnight at 4°C, followed by incubation with Alexa Fluor 488-conjugated goat anti-mouse IgG (1:2000; Thermo Fisher Scientific, A32723) or Cy3-labeled Goat Anti-Rat IgG(H+L) (1:500; Beyotime, A0507) for 1 h at RT in the dark. Nuclei were counterstained with fluorescent brightener (Sigma, MKCL1227) for 5 min. Coverslips were mounted with antifade medium (Solabio) and imaged using a Zeiss LSM 980 confocal microscope (63×oil objective) with Z-stack acquisition (0.5 μm intervals). Maximum intensity projections were generated using ZEN Blue software. Fluorescence intensity was quantified in ImageJ with consistent background subtraction and thresholding across samples. Primary antibodies: anti-EnP1 mouse 1:200,anti-SWP1b Rat 1:200.

### Sequence and Structural Homology Analysis and Secondary Structure Analysis

Sequence similarity searches were performed using BLASTp and PSI-BLAST (https://blast.ncbi.nlm.nih.gov/Blast.cgi) against the clustered non-redundant (nr) database^54^. Significant hits (*E* ≤ 1 × 10^−5^) were used to assess the taxonomic distribution of homologs.

Structural homology was evaluated using HHpred (https://toolkit.tuebingen.mpg.de/tools/hhpred) against PDB-derived databases^55^. Reciprocal BLASTp searches were performed for candidate remote homologs.

Secondary structure analysis was performed with PSIPRED (https://bioinf.cs.ucl.ac.uk/psipred/). PSIPRED 4.0 was used to predict the secondary structure and DISOPRED3 was used to predict disordered regions using a cutoff of 0.5^56,57^.

### AlphaFold3 Modeling

Individual proteins and the Swp1b-EnP1 multimer were modeled with the AlphaFold3 server (https://alphafoldserver.com/) with standard settings^4^.

### Integrative Structural Modeling

After crosslinks were extracted with RRI_FDR.R intralinks were extracted with getAlphaLinkFile.R. All intralinks with an FDR ≤ 0.05 were included. Initial structural models of Swp1b and EnP1 were generated using AlphaLink, a modified version of OpenFold that incorporates experimental distance restraints into structural prediction ^26^. Intralinks were provided as distance distributions generated by AlphaLink. 30Å was provided as the upper bound restraint. For each protein the relaxed output model was selected for downstream analysis.

The EnP1 model was modified so it was labeled as chain “B” and transformed 150Å in the x direction by modify_pdb.R to clearly separate them from one other. The two protein monomers were used as templates and brought together using MODELLER (https://salilab.org/modeller/), incorporating all intralinks and interlinks with FDR ≤ 0.05 as 30Å upper-bound distance restraints with a standard deviation of 0.1 by redefining the AutoModel.special_restraints() routine as described in the MODELLER manual^36^.

All scripts used and an example .pir alignment file are provided in Supplemental Materials.

Crosslinks are mapped onto the final model and their distances are measured with calculateMultimerDistance.py by creating distance objects in PyMOL^58^. RMS Z-scores were obtained from the WHATCHECK tool (https://bio.tools/what_check) with default settings^41^. QMEAN (available via SWISSMODEL) (https://swissmodel.expasy.org/qmean/) score generated the confidence for each residue^42^. VMD was utilized to analyze the number of hydrogen bonds and salt bridges (http://www.ks.uiuc.edu/Research/vmd/)^40^.

## Supporting information

Scripts and Example Files

## Acknowledgements

The Sidoli lab gratefully acknowledges for funding the Hevolution Foundation (AFAR), the ERCM-CFAR Center for AIDS Research, the NIAID for the PROVIDENT grant (1U19AI181977), the Einstein-Mount Sinai Diabetes center, and the NIH Office of the Director (S10OD030286). E.W. is supported by NIH grant R01AI311965.

## SUPPLEMENTAL FIGURES, DATA, SCRIPTS, AND FILES

<u>Supplemental Figures 1-6</u>

<u>Supplemental Data 1</u>

List of Swp1b-EnP1 crosslinks

<u>Supplemental Data 2</u>

List of independent set of crosslinks

<u>RRI_FDR.R</u>

Extract crosslink data from Annika2.0.

<u>getAlphaLinkFile.R</u>

Extract intralinks for proteins of interest and put in AlphaLink format.

<u>modify_pdb.R</u>

Modify pdb file for MODELLER format.

<u>xl-for-modeller.R</u>

Export crosslinks and intralinks in MODELLER format.

<u>Swp1b-EnP1.pir</u>

Example .pir file for bringing together two chains in MODELLER.

<u>model_w_restraints.py</u>

Example file to run MODELLER.

<u>calculateMultimerDistance.py</u>

Map and record crosslink distances in PyMOL.

## References

1. Han, B. & Weiss, L. M. Microsporidia: Obligate Intracellular Pathogens within the Fungal Kingdom. Microbiol Spectr 5, 10.1128/microbiolspec.FUNK-0018–2016 (2017).

2. Keeling, P. J. & Fast, N. M. Microsporidia: biology and evolution of highly reduced intracellular parasites. Annu Rev Microbiol 56, 93–116 (2002).

3. Nakjang, S. et al. Reduction and Expansion in Microsporidian Genome Evolution: New Insights from Comparative Genomics. Genome Biol Evol 5, 2285–2303 (2013).

4. Abramson, J. et al. Accurate structure prediction of biomolecular interactions with AlphaFold 3. Nature 630, 493–500 (2024).

5. Baek, M. et al. Accurate prediction of protein structures and interactions using a three-track neural network. Science 373, 871–876 (2021).

6. Lee, J.-W. et al. DeepFold: enhancing protein structure prediction through optimized loss functions, improved template features, and re-optimized energy function. Bioinformatics 39, btad712 (2023).

7. Wu, R. et al. High-resolution de novo structure prediction from primary sequence. 2022.07.21.500999 Preprint at 10.1101/2022.07.21.500999 (2022).

8. Evolutionary-scale prediction of atomic-level protein structure with a language model | Science. https://www.science.org/doi/10.1126/science.ade2574.

9. Varadi, M. et al. AlphaFold Protein Structure Database in 2024: providing structure coverage for over 214 million protein sequences. Nucleic Acids Res 52, D368–D375 (2023).

10. Sarti, E. & Cazals, F. Fold or flop: quality assessment of AlphaFold predictions on whole proteomes. 2025.12.19.695427 Preprint at 10.64898/2025.12.19.695427 (2026).

11. Jing, X., Wu, F., Luo, X. & Xu, J. Single-sequence protein structure prediction by integrating protein language models. Proc Natl Acad Sci U S A 121, e2308788121.

12. Akdel, M. et al. A structural biology community assessment of AlphaFold2 applications. Nat Struct Mol Biol 29, 1056–1067 (2022).

13. Ahdritz, G. et al. OpenFold: retraining AlphaFold2 yields new insights into its learning mechanisms and capacity for generalization. Nat Methods 21, 1514–1524 (2024).

14. Li, M. Q. C. et al. Advantages and Limitations of AlphaFold in Structural Biology: Insights from Recent Studies. Protein J 45, 22–38 (2026).

15. Park, S., Myung, S. & Baek, M. Advancing protein structure prediction beyond AlphaFold2. Current Opinion in Structural Biology 90, 102985 (2025).

16. Meng, Q., Guo, F. & Tang, J. Improved structure-related prediction for insufficient homologous proteins using MSA enhancement and pre-trained language model. Brief Bioinform 24, bbad217 (2023).

17. Dunker, A. K. et al. Intrinsically disordered protein. Journal of Molecular Graphics and Modelling 19, 26–59 (2001).

18. Barchet, C. et al. Focused classifications and refinements in high-resolution single particle cryo-EM analysis. Journal of Structural Biology 215, 108015 (2023).

19. Noone, D. P. et al. PTX3 structure determination using a hybrid cryoelectron microscopy and AlphaFold approach offers insights into ligand binding and complement activation. Proceedings of the National Academy of Sciences 119, e2208144119 (2022).

20. Alshammari, M., He, J. & Wriggers, W. Flexible fitting of AlphaFold2-predicted models to cryo-EM density maps using elastic network models: a methodical affirmation. Bioinformatics Advances 5, vbae181 (2025).

21. Barbarin-Bocahu, I. & Graille, M. The X-ray crystallography phase problem solved thanks to AlphaFold and RoseTTAFold models: a case-study report. Acta Cryst D 78, 517–531 (2022).

22. Gyawali, R., Dhakal, A. & Cheng, J. Multimodal deep learning integration of cryo-EM and AlphaFold3 for high-accuracy protein structure determination. Commun Chem 8, 320 (2025).

23. Klykov, O. et al. Efficient and robust proteome-wide approaches for cross-linking mass spectrometry. Nat Protoc 13, 2964–2990 (2018).

24. Liu, S. et al. Assisting and accelerating NMR assignment with restrained structure prediction. Commun Biol 8, 1067 (2025).

25. Ziemianowicz, D. S. et al. IMProv: A Resource for Cross-link-Driven Structure Modeling that Accommodates Protein Dynamics. Mol Cell Proteomics 20, 100139 (2021).

26. Stahl, K., Graziadei, A., Dau, T., Brock, O. & Rappsilber, J. Protein structure prediction with in-cell photo-crosslinking mass spectrometry and deep learning. Nat Biotechnol 41, 1810–1819 (2023).

27. Stahl, K. et al. Modelling protein complexes with crosslinking mass spectrometry and deep learning. Nat Commun 15, 7866 (2024).

28. Zhang, Y. et al. Distance-AF improves predicted protein structure models by AlphaFold2 with user-specified distance constraints. Commun Biol 8, 1392 (2025).

29. Xie, Y. et al. Integrating diverse experimental information to assist protein complex structure prediction by GRASP. Nat Methods 22, 2362–2374 (2025).

30. Honorato, R. V. et al. The HADDOCK2.4 web server for integrative modeling of biomolecular complexes. Nat Protoc 19, 3219–3241 (2024).

31. Gong, Z.Ye, S.-X. & Tang, C. Tightening the Crosslinking Distance Restraints for Better Resolution of Protein Structure and Dynamics. Structure 28, 1160-1167.e3 (2020).

32. Han, B., Pan, G. & Weiss, L. M. Microsporidiosis in Humans. Clin Microbiol Rev 34, e00010–20.

33. Peuvel-Fanget, I. et al. EnP1 and EnP2, two proteins associated with the Encephalitozoon cuniculi endospore, the chitin-rich inner layer of the microsporidian spore wall. International Journal for Parasitology 36, 309–318 (2006).

34. Guan, J. et al. Microsporidian EnP1 alters host cell H2B monoubiquitination and prevents ferroptosis facilitating microsporidia survival. Proc Natl Acad Sci U S A 121, e2400657121.

35. Polonais, V. et al. The Human Microsporidian Encephalitozoon hellem Synthesizes Two Spore Wall Polymorphic Proteins Useful for Epidemiological Studies. Infect Immun 78, 2221–2230 (2010).

36. Šali, A. & Blundell, T. L. Comparative Protein Modelling by Satisfaction of Spatial Restraints. Journal of Molecular Biology 234, 779–815 (1993).

37. Burke, A. M. et al. Synthesis of two new enrichable and MS-cleavable cross-linkers to define protein–protein interactions by mass spectrometry. Org. Biomol. Chem. 13, 5030–5037 (2015).

38. Merkley, E. D. et al. Distance restraints from crosslinking mass spectrometry: Mining a molecular dynamics simulation database to evaluate lysine–lysine distances. Protein Sci 23, 747–759 (2014).

39. O’Reilly, F. J. & Rappsilber, J. Cross-linking mass spectrometry: methods and applications in structural, molecular and systems biology. Nat Struct Mol Biol 25, 1000–1008 (2018).

40. Humphrey, W., Dalke, A. & Schulten, K. VMD: Visual molecular dynamics. Journal of Molecular Graphics 14, 33–38 (1996).

41. Hooft, R., Vriend, G., Sander, C. & Abola, E. Errors in protein structures. Nature 381, 272 (1996).

42. Benkert, P., Biasini, M. & Schwede, T. Toward the estimation of the absolute quality of individual protein structure models. Bioinformatics 27, 343–350 (2011).

43. Fernandez-Fuentes, N., Dybas, J. M. & Fiser, A. Structural Characteristics of Novel Protein Folds. PLOS Computational Biology 6, e1000750 (2010).

44. Bigliardi, E. et al. Microsporidian Spore Wall: Ultrastructural Findings on Encephalitozoon hellem Exospore. Journal of Eukaryotic Microbiology 43, 181–186 (1996).

45. Tomita, T. et al. Mapping a Toxoplasma gondii interactome by crosslinking mass spectrometry and machine learning. mBio 16, e02159–25.

46. Matzinger, M., Kandioller, W., Doppler, P., Heiss, E. H. & Mechtler, K. Fast and Highly Efficient Affinity Enrichment of Azide-A-DSBSO Cross-Linked Peptides. J Proteome Res 19, 2071–2079 (2020).

47. Stieger, C. E., Doppler, P. & Mechtler, K. Optimized Fragmentation Improves the Identification of Peptides Cross-Linked by MS-Cleavable Reagents. J. Proteome Res. 18, 1363–1370 (2019).

48. Klössel, S. et al. Yeast TLDc domain proteins regulate assembly state and subcellular localization of the V-ATPase. EMBO J 43, 1870–1897 (2024).

49. Schnirch, L. et al. Expanding the Depth and Sensitivity of Cross-Link Identification by Differential Ion Mobility Using High-Field Asymmetric Waveform Ion Mobility Spectrometry. Anal. Chem. 92, 10495–10503 (2020).

50. Amos, B. et al. VEuPathDB: the eukaryotic pathogen, vector and host bioinformatics resource center. Nucleic Acids Res 50, D898–D911 (2021).

51. Birklbauer, M. J., Matzinger, M., Müller, F., Mechtler, K. & Dorfer, V. MS Annika 2.0 Identifies Cross-Linked Peptides in MS2–MS3-Based Workflows at High Sensitivity and Specificity. J Proteome Res 22, 3009–3021 (2023).

52. Lenz, S. et al. Reliable identification of protein-protein interactions by crosslinking mass spectrometry. Nat Commun 12, 3564 (2021).

53. Combe, C. W., Graham, M., Kolbowski, L., Fischer, L. & Rappsilber, J. xiVIEW: Visualisation of Crosslinking Mass Spectrometry Data. Journal of Molecular Biology 436, 168656 (2024).

54. Altschul, S. F. et al. Gapped BLAST and PSI-BLAST: a new generation of protein database search programs. Nucleic Acids Res 25, 3389–3402 (1997).

55. Söding, J., Biegert, A. & Lupas, A. N. The HHpred interactive server for protein homology detection and structure prediction. Nucleic Acids Res 33, W244–W248 (2005).

56. McGuffin, L. J., Bryson, K. & Jones, D. T. The PSIPRED protein structure prediction server. Bioinformatics 16, 404–405 (2000).

57. Jones, D. T. & Cozzetto, D. DISOPRED3: precise disordered region predictions with annotated protein-binding activity. Bioinformatics 31, 857–863 (2015).

58. Schrödinger, LLC. The PyMOL Molecular Graphics System, Version 3.1. (2015).

