## Supplemental Figures for "XL-MS–Guided Structure Prediction of Disordered *Encephalitozoon hellem* Proteins"

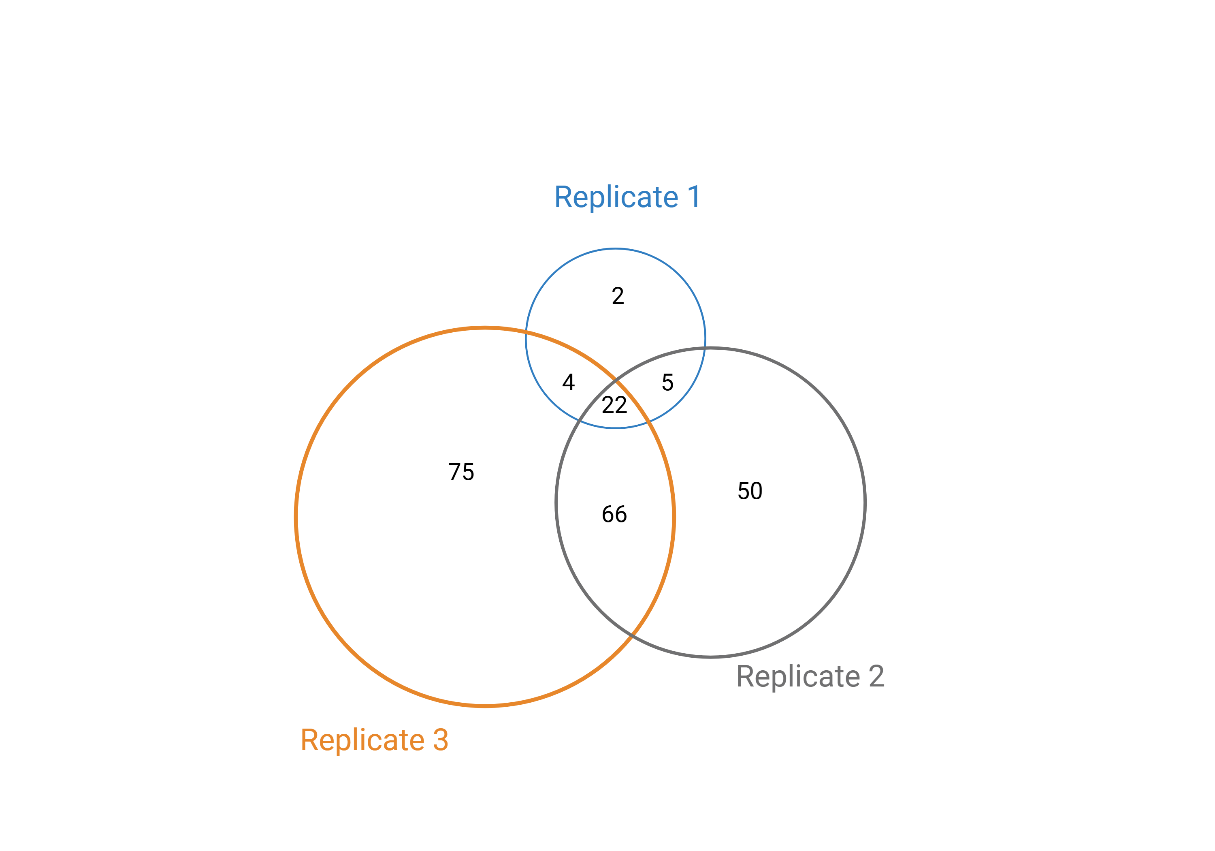

**Supplemental Figure 1|** Number of crosslinks (intralinks and interlinks) found in each independent replicate that contributed to the Swp1b-EnP1 model. Created with Biorender.com.

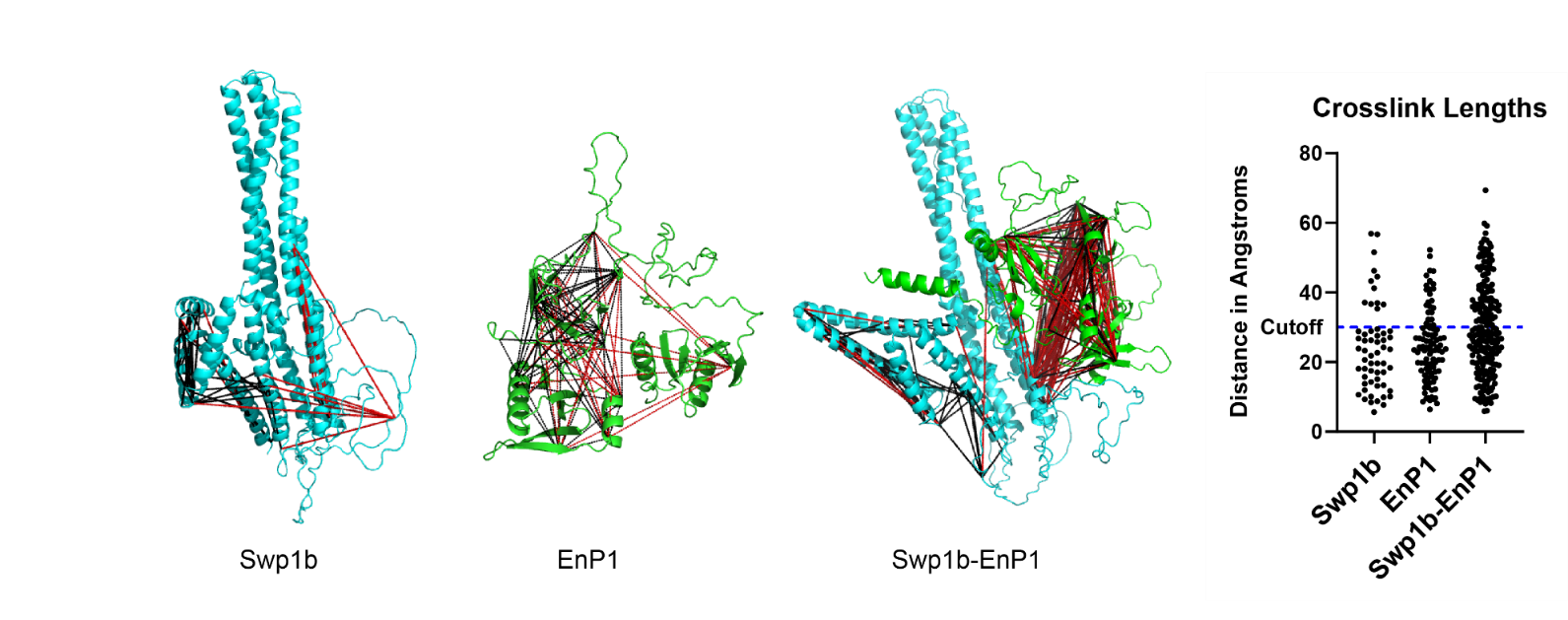

**Supplemental Figure 2**| XL-MS crosslinks mapped onto Swp1b, EnP1, and Swp1b-EnP1 structures generated by ESMFold. Links below the 30Å cutoff are colored black, links above the 30Å are colored red. 128/224 crosslinks (57%) fall below the 30Å cutoff in the Swp1b-EnP1 complex.

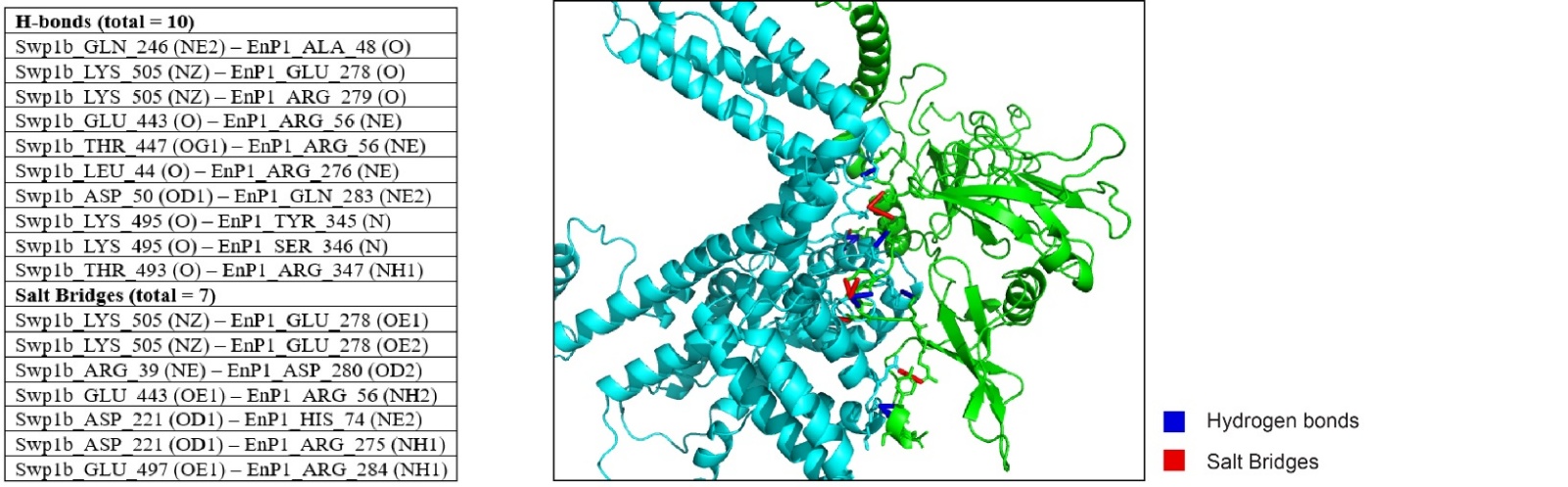

**Supplemental Figure 3|Analysis of Hydrogen bonds and Salt Bridges.** List of hydrogen bonds (H-bonds) and salt bridges between Swp1b and EnP1 atoms (left) and mapped onto the model (right).

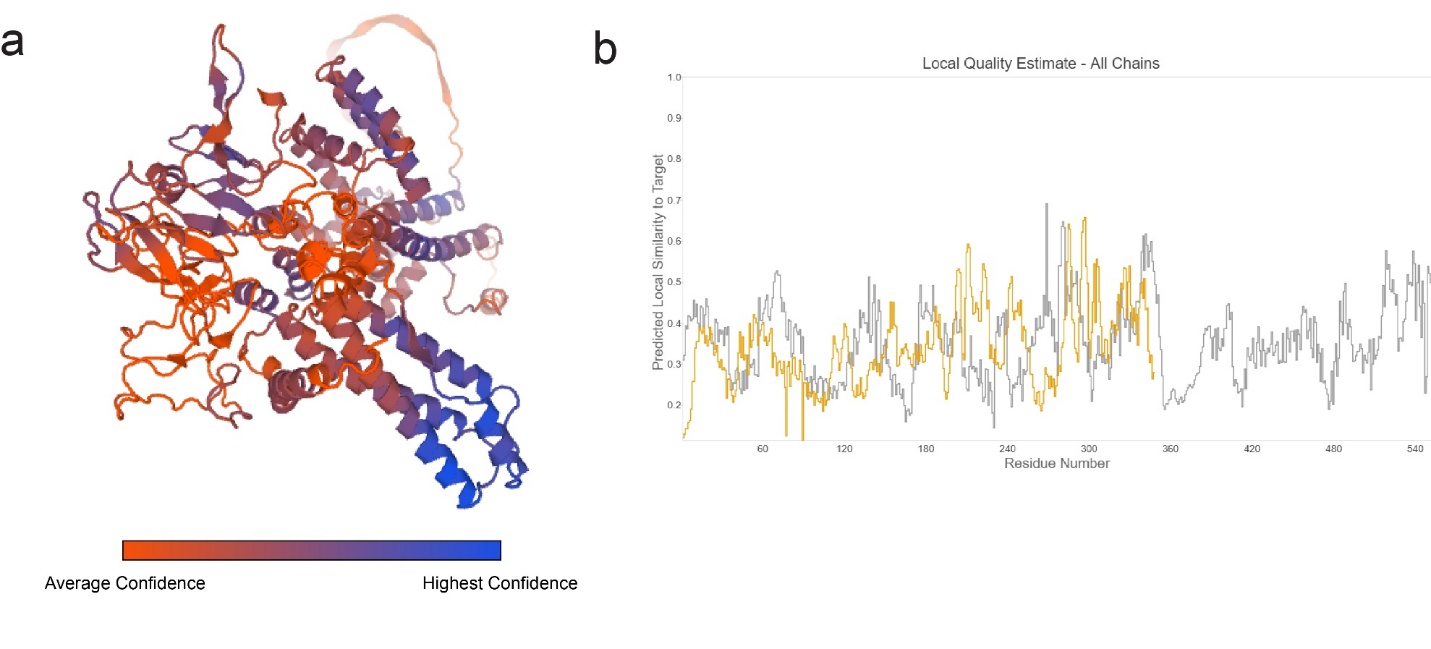

**Supplemental Figure 4| Quality analysis of Swp1b-EnP1 model a.** Per-residue confidence as determined by QMEAN. **b.** Local quality estimate as determined by WHATCHECK.

| Plasmids | Selective Media | Expected Result if Positive | Swp1b-FL  EnP1-FL | Swp1b^32-353^  EnP1^141-348^ | (GGGGS)_3_-Swp1b-FL  (GGGGS)_3_-EnP1-FL | Swp1b^20-60^  EnP1^141-215^ |
| --- | --- | --- | --- | --- | --- | --- |
| **BD-52**  **AD-T**  **(Positive Control)** | -Leu/-Trp | + | + | + | + | + |
|  | -Leu/-Trp/XaG/Aba | + blue | + blue | + blue | + blue | + blue |
| **BD-Lam**  **AD-T**  **(Negative Control)** | -Leu/-Trp | + | + | + | + | + |
|  | -Leu/-Trp/XaG/Aba | - | - | - | - | - |
| **BD-Swp1b (test for autoactivation)** | -Trp | + | + | + | + | + |
|  | -Trp/XaG | + white or blue | + blue | + blue | + blue | + blue |
|  | -Trp/XaG/Aba | - | - | - | - | - |
| **BD Empty (test for bait toxicity)** | -Trp | + | + | + | + | + |
| **BD Empty**  **AD-EnP1 (test for false positive)** | -Leu/-Trp/XaG | + white | + blue | + blue | + white | + white |
|  | -Ade/-His/-Leu/-Trp/XaG/AbA | - | - | - | - | - |
| **BD-Swp1b**  **AD-EnP1** | -Leu/-Trp/XaG | + blue | + blue | + blue | + blue | + blue |
|  | -Ade/-His/-Leu/-Trp/XaG/Aba | + blue | - | - | - | - |

**Supplemental Figure 5| Yeast-2-hybrid analysis of Swp1b and EnP1 interaction**. Four versions of Swp1b and EnP1 were tested. The full length of each protein (Swp1b-FL and EnP1-FL), approximately half of each protein (Swp1b^32-353^ and EnP1^141-348^), small pieces of the protein predicted to interact (Swp1b^20-60^ and EnP1^141-215^, and the full-length proteins with a flexible linker added to the N terminus ((GGGGS)_3_-Swp1b-FL and (GGGGS)_3_-EnP1-FL). All controls (first 10 rows) worked as expected. No interaction between Swp1b and EnP1 was detected in any of the four versions, as they failed to grow on -Ade/-His/-Leu/-Trp/XaG/Aba selective media.

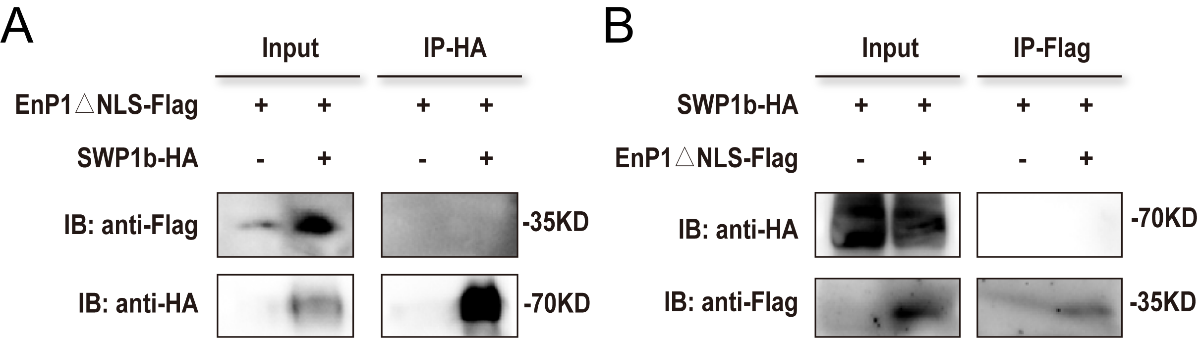

**Supplemental Figure 6| Co-immunoprecipitation does not reveal an interaction between EnP1ΔNLS-Flag and SWP1b-HA** Co-immunoprecipitation (Co-IP) assays of the interaction between EnP1ΔNLS-Flag and SWP1b-HA. **a.** Lysates from cells co-expressing EnP1ΔNLS-Flag and SWP1b-HA were immunoprecipitated with anti-HA magnetic beads (IP-HA). Inputs and immunoprecipitates were immunoblotted (IB) with anti-Flag (upper panel) and anti-HA (lower panel) antibodies. **b.** Reciprocal co-IP assays using anti-Flag magnetic beads (IP-Flag). IB was performed with anti-HA (upper panel) and anti-Flag (lower panel) antibodies. Molecular weights (kDa) are indicated on the right. Experiments were independently repeated using HA magnetic beads and Flag magnetic beads, respectively.
