## Supplementary material for "XL-MS–Guided Structure Prediction of Disordered *Encephalitozoon hellem* Proteins": Scripts and Example Files: calculateMultimerDistance.py

```
# -*- coding: utf-8 -*-
"""
Created on Tue Aug 26 15:07:58 2025

@author: Elizabeth Weyer
"""

#Manage working directory in Spyder (not necessary if you aren't making changes to this code)
    #import os
    #os.getcwd()
    #os.chdir("C:/Users/lmwla/OneDrive - Montefiore Medicine/Documents/Swp1b-EnP1_Modelling")

#### READ BEFORE USING ####

#Install pandas in pymol environment. Only need to do this once.
#Find pymol's python environment:
    #import sys
    #print(sys.exectuable)
#copy path and open command prompt in your computer. This is for windows
#C:\path -m pip install pandas
#Replace path with the environment path

#Instructions for running in PyMOL:

#Set your working directory. Make sure all files are in this directory
#Example: cd C:\Users\lmwla\OneDrive - Montefiore Medicine\Documents\Swp1b-EnP1_Modelling

#Make sure your input file is in the right format. It should be in the format you used to run Haddock (chain1, chain2, residue1, residue2).

#Run python file. Make sure your working directory in pymol is the same as this file and the other files you need
#Example: run C:\Users\lmwla\OneDrive - Montefiore Medicine\Documents\Swp1b-EnP1_Modelling\Haddock_New_Models\calculateDistance.py

#Next run the program, input the arguments. They all need to be in quotes to work
#Example: calculateMultimerDistance("All-XL.csv", "Swp1b-EnP1", "30", "Swp1b-EnP1.csv")
    #First argument = input file you used for Haddock
    #Second argument = name of the model in PyMOL
    #Third argument = Distance cutoff
    #Fourth argument = Name of your file to output the distances

#### END OF INSTRUCTIONS ####

def calculateMultimerDistance(file, model_name, distance, out_file):

    #Import packages needed
    import pandas as pd
    from pymol import cmd
    
    #Rename arguments
    file_name = str(file)
    mol1 = str(model_name)
    dist = str(distance)
    out_file_name = str(out_file)

    #Read in table from Haddock format and clean table
    df = pd.read_csv(file_name, sep=",")
    df['residue1'] = df['residue1'].astype(int)
    df['residue2'] = df['residue2'].astype(int)

    #Open a file to write to. Change name.
    f=open(out_file_name, 'w')

    #Go through the table, create distance object, write distance to file
    for index, row in df.iterrows():
        name = dist + "A_d" + str(index+1)
        res1 = mol1 + " and chain " + row["chain1"] + " and resi " + str(int(row["residue1"])) + " and name CA"
        res2 = mol1 + " and chain " + row["chain2"] + " and resi " + str(int(row["residue2"])) + " and name CA"
        dst=cmd.distance(name, res1, res2)
        #f.write(res1)
        #f.write("%8.3f\n"%dst)
        f.write(f"{name} {dst:.3f}\n")

    #Close file
    f.close()

#Add so pymol can read this file
cmd.extend( "calculateMultimerDistance", calculateMultimerDistance );

#Clean up image - you can run these in PyMOL command line
#hide labels, 30A_d*
#color black, 30A_d*
```
