## Supplementary material for "XL-MS–Guided Structure Prediction of Disordered *Encephalitozoon hellem* Proteins": Scripts and Example Files: getAlphaLinkFile.R

```
library(tidyverse)

#read in file. Make sure to set your working directory.
df <- read.csv("AlphaLink_RRI.csv", sep = "\t")

#get protein you want (hard code this in)
accession = "EHEL_101700"

#Only select rows for the protein of interest
selected_rows <- subset(df, grepl(accession, Accession.A))

#Change "High" to 0.01 and "Medium" to 0.05
selected_rows$Confidence[selected_rows$Confidence == "High"] <- 0.01
selected_rows$Confidence[selected_rows$Confidence == "Medium"] <- 0.05

#Remove any rows where the difference in amino acids is <= 3. This means that it is likely a link between the same peptide in two proteins.
#If you want these links to model a dimer they are output to moved_rows and you can write these out into a file.
selected_rows$In.protein.A <- as.numeric(selected_rows$In.protein.A)
selected_rows$In.protein.B <- as.numeric(selected_rows$In.protein.B)
selected_rows$distance <- selected_rows$In.protein.A - selected_rows$In.protein.B
kept_rows <- selected_rows[abs(selected_rows$distance) > 3, ]
moved_rows <- selected_rows[abs(selected_rows$distance) <= 3, ]

#Export in Alphalink format
ExportData <- kept_rows %>% select(
  In.protein.A,
  In.protein.B,
  Confidence
)
#Export for AlphaLink. Change file name
write.table(ExportData, "Swp1b_IntraLinks.csv", row.names = FALSE, col.names = FALSE, sep = "\t", quote = FALSE)
```
