## Supplementary material for "XL-MS–Guided Structure Prediction of Disordered *Encephalitozoon hellem* Proteins": Scripts and Example Files: model_w_restraints.py

```
# Addition of restraints to the default ones
from modeller import *
from modeller.automodel import *    # Load the AutoModel class
from modeller import soap_pp

log.verbose()
env = Environ()

# directories for input atom files
env.io.atom_files_directory = ['.', '../atom_files']

class MyModel(AutoModel):
    def special_restraints(self, aln):
        rsr = self.restraints
        at = self.atoms
        #Use xl-for-modeller.R to generate your restraints in the correct format, then paste them here.
        '''
        rsr.add(forms.upper_bound(group=physical.xy_distance,
                                    feature=features.Distance(at['CA:217:A'],
                                                                at['CA:703:B']),
                                        mean=30, stdev=0.1))
        '''


        
a = MyModel(env,
            alnfile  = 'Swp1b-EnP1.pir',     # alignment filename
            knowns   = ('Swp1b', 'EnP1-translated'),   # codes of the templates
            sequence = 'Swp1b_EnP1',         # code of the target
            assess_methods=(assess.DOPE, soap_pp.Assessor(), assess.GA341))     

a.starting_model= 1                 # index of the first model
a.ending_model  = 1                # index of the last model
# (determines how many models to calculate)

#Thourough VTFM optimization
a.library_schedule = autosched.slow
a.max_var_interations = 300

#thorough MD optimization
a.md_level = refine.very_slow

a.make()                            # do comparative modeling
```
