## Supplementary material for "XL-MS–Guided Structure Prediction of Disordered *Encephalitozoon hellem* Proteins": Scripts and Example Files: modify_pdb.R

```
#to get headers:
#str(pdb)
#head(pdb)

library(Rpdb)

pdb <- read.pdb("EnP1.pdb")

pdb$ATOM$chain <- "B"
pdb$HETATM$chain <- "B"

pdb$ATOM$x <- pdb$ATOM$x + 150
pdb$HETATM$x <- pdb$HETATM$x + 150

write.pdb(pdb, file = "EnP1-translated.pdb")
```
