## Supplementary material for "XL-MS–Guided Structure Prediction of Disordered *Encephalitozoon hellem* Proteins": Scripts and Example Files: RRI_FDR.R

```
library(tidyverse)

########################################
#Clean data
########################################

#Load the clean full XLMS dataframe 

#read the "crosslinks" (= peptide pairs level) results from the Annika 2.0 results
rawdf <- read.table("DSBSO_Crosslinks.txt", header = TRUE, sep = "\t")

#removes suffix on protein names and isoform designation
rawdf <- rawdf %>%
  mutate(across(where(is.character), ~str_replace_all(., "-t26_1-p1", ""))) %>%
  mutate(across(where(is.character), ~str_replace_all(., "\\.P\\d{2,4}", "")))

#make the descriptions easy to read
# Define a function to clean specified columns with specific patterns, prefix removal, and custom string removals
clean_gene_product <- function(data, columns) {
  # Apply multiple transformations within mutate across the specified columns
  data %>% 
    mutate(across(all_of(columns), ~if_else(
      str_starts(., "transcript"),  # Check if starts with "transcript"
      str_split(., "\\|", simplify = TRUE)[, 4],  # Split and select the 4th element if condition is met
      .  # Keep the original value if condition is not met
    ))) %>%
    mutate(across(all_of(columns), ~str_remove(., "^\\s*gene_product=")))  # Remove the "gene_product=" prefix
} 
# apply that funciton to df
rawdf <- clean_gene_product(rawdf, c("Protein.Descriptions.A", "Protein.Descriptions.B"))


#remove ambiguous peptides matching multiple proteins
rawdf2 <- rawdf %>% filter(!str_detect(Accession.A, ";") & !str_detect(Accession.B, ";"))

#RRI CONDENSATION
RRIdf <- rawdf2 %>%
  #add Decoy.Pair column with TT/TD/DD for FDR calculation 
  rowwise() %>%
  mutate(
    Decoy.Pair = case_when(
      # both Accession.A and Accession.B are empty or NA
      (is.na(Accession.A) | Accession.A == "") &
        (is.na(Accession.B) | Accession.B == "") ~ "DD",
      
      # exactly one of them is empty/NA → “TD”
      (is.na(Accession.A) | Accession.A == "") &
        !(is.na(Accession.B) | Accession.B == "") ~ "TD",
      
      !(is.na(Accession.A) | Accession.A == "") &
        (is.na(Accession.B) | Accession.B == "") ~ "TD",
      
      # if neither is empty/NA → “TT”
      TRUE ~ "TT"
    )
  ) %>%
  ungroup() %>%
  #Give unique decoy ID so that they don't collapse into one DD
  # 1) Tag every row/column for “is empty?”
  mutate(
    is_empty_A = is.na(Accession.A) | Accession.A == "",
    is_empty_B = is.na(Accession.B) | Accession.B == ""
  ) %>%
  # 2) Compute total number of empty cells across A+B (same for every row)
  mutate(total_empty = sum(is_empty_A) + sum(is_empty_B)) %>%
  
  # 3) Generate exactly `total_empty` unique 6-digit random numbers, 
  #    zero-pad them, and prepend “DECOY_”; store as a list-column `all_decoys`
  mutate(all_decoys = list(
    str_c(
      "DECOY_",
      str_pad(
        sample(0:999999, size = total_empty[1], replace = FALSE),
        width = 6,
        pad = "0"
      )
    )
  )) %>%
  
  # 4) Assign a row-wise index for each empty spot in A and B:
  #    - “index_A” runs 1, 2, … across all rows where is_empty_A == TRUE
  #    - “index_B” runs (1 + #empty_A), (2 + #empty_A), … across rows where is_empty_B == TRUE
  mutate(
    # Within all rows, do a running count of TRUEs in is_empty_A
    index_A = if_else(is_empty_A, cumsum(is_empty_A), NA_integer_),
    # For B, offset its cumsum by sum(is_empty_A) so IDs don’t collide
    index_B = if_else(is_empty_B, cumsum(is_empty_B) + sum(is_empty_A), NA_integer_)
  ) %>%
  
  # 5) Now pull from the “all_decoys” vector to fill each empty cell:
  mutate(
    Accession.A = if_else(
      is_empty_A,
      # pick the index-th decoy from all_decoys[[1]]
      all_decoys[[1]][index_A],
      Accession.A
    ),
    Accession.B = if_else(
      is_empty_B,
      all_decoys[[1]][index_B],
      Accession.B
    )
  ) %>%
  
  # 6) Drop all helper columns
  select(-is_empty_A, -is_empty_B, -total_empty, -all_decoys, -index_A, -index_B) %>%
  
  #Add 0 to the decoy's residue number  
  mutate(across(
    c(In.protein.A, In.protein.B),
    ~ if_else(is.na(.) | . == "", "0", .)
  )) %>%
  mutate(across(c(In.protein.A, In.protein.B), as.numeric)) %>%
  
  # --- (a) Create Residue‐to‐Residue Interaction (RRI) label --- #
  rowwise() %>%
  mutate(
    RRI = {
      # build the two “protein:position” strings on the fly
      R1 <- str_c(Accession.A, In.protein.A, sep = ":")
      R2 <- str_c(Accession.B, In.protein.B, sep = ":")
      # sort them and paste with “-”
      paste(sort(c(R1, R2)), collapse = "-")
    }
  ) %>%
  ungroup() %>%
  
  # --- (b) Aggregate all peptide‐level entries sharing the same RRI --- #
  group_by(RRI) %>%
  mutate(
    Number.of.CSMs = sum(Number.of.CSMs),  # replace by group sum
    RRI.Score = sqrt(sum(Best.CSM.Score^2)) #Euclidean-norm of Best.CSM.Score
  ) %>%
  slice_max(order_by = Best.CSM.Score, n = 1, with_ties = FALSE) %>%
  ungroup() %>%
  
  # --- (c) Create Protein‐to‐Protein Interaction (PPI) label --- #
  rowwise() %>%
  mutate(
    PPI = paste(sort(c(Accession.A, Accession.B)), collapse = "-")
  ) %>%
  ungroup()


########################################
#CALCULATE FDR AT RRI LEVEL
########################################

RRIfdrcalc <- RRIdf %>%
  filter(Crosslink.Type == "Inter") %>%
  #1) arrange so that highest score to lowest score
  arrange(desc(RRI.Score)) %>%
  #2) add cumulative counts
  mutate(
    CumTD = cumsum(Decoy.Pair == "TD"),
    CumDD = cumsum(Decoy.Pair == "DD"),
    CumTT = cumsum(Decoy.Pair == "TT"),
    RRI_FDR = (CumTD - CumDD)/CumTT)

desired_FDRs <- c(0.01, 0.05, 0.10)

cutoff_table_RRI <- tibble(FDR_target = desired_FDRs) %>%
  mutate(
    cutoff_score = map_dbl(FDR_target, ~ {
      tmp <- RRIfdrcalc %>%
        filter(RRI_FDR <= .x)
      if (nrow(tmp) == 0) {
        # no row ever achieves FDR <= .x → return NA
        NA_real_
      } else {
        # take the bottommost (lowest‐score) row among those that pass
        tmp %>%
          slice_tail(n = 1) %>%
          pull(RRI.Score)
      }
    })
  )

########################################
# Get rid of decoy
########################################

RRI_clean <- RRIdf %>% filter(Decoy == "False") %>%
  # Modify 'Confidence' based on new conditions
  arrange(desc(RRI.Score)) %>%
  mutate(Confidence_RRI = case_when(
    Crosslink.Type == "Intra" ~ Confidence, # Keep original values from the column Confidence for intralinks
    RRI.Score > cutoff_table_RRI %>% filter(FDR_target == 0.01) %>% pull(cutoff_score) ~ "High", # FDR <1%
    RRI.Score > cutoff_table_RRI %>% filter(FDR_target == 0.05) %>% pull(cutoff_score) ~ "Medium", # FDR <5%
    RRI.Score <= cutoff_table_RRI %>% filter(FDR_target == 0.10) %>% pull(cutoff_score) ~ "Low", # FDR <10%
    TRUE ~ Confidence # Default to keep original in all other cases
  ))


########################################
#Get Medium and High IntraLinks for AlphaLink
########################################

alphaLink_df <- RRI_clean %>% 
  filter(Confidence != "Low") %>%  
  filter(Crosslink.Type == "Intra")

ExportData_AlphaLink <- alphaLink_df %>% select(
  Accession.A,
  Accession.B,
  Protein.Descriptions.A,
  Protein.Descriptions.B,
  In.protein.A,
  In.protein.B,
  Confidence
)
write.table(ExportData_AlphaLink, "AlphaLink_RRI.csv", sep = "\t", row.names = FALSE, quote = FALSE)

########################################
#Get Medium and High InterLinks for Modelling
########################################

modelling_df <- RRI_clean %>% 
  filter(Confidence != "Low") %>%  
  filter(Crosslink.Type == "Inter")

ExportData_Modelling <- modelling_df %>% select(
  Accession.A,
  Accession.B,
  Protein.Descriptions.A,
  Protein.Descriptions.B,
  In.protein.A,
  In.protein.B,
  Confidence
)

write.table(ExportData_Modelling, "Modelling_RRI.csv", sep = "\t", row.names = FALSE, quote = FALSE)
```
