## Supplementary material for "XL-MS–Guided Structure Prediction of Disordered *Encephalitozoon hellem* Proteins": Scripts and Example Files: xl-for-modeller.R

```
#We are going to load in all of our crosslinks and intralinks and output them to copy paste in the python file.This will also make a file that can
#be used with CompareMultimerDistance.py to map links onto the structure in PyMOL.

#Read in Modeeling_RRI file
df <- read.table("Modelling_RRI.csv", sep ="\t", header = TRUE)                                    

#read in alphalink file for chain A
df_A <- read.table("Swp1b_IntraLinks.csv", sep="\t")                          

#read in alphalink file for chain B
df_B <- read.table("EnP1_IntraLinks.csv", sep="\t")                        

#Input your distance cutoff
distance_cutoff <- 30                                                         

#Input your standard deviation (usually 0.1)
std_dev <- 0.1

#This is the length of chain A that will be added to chain B
chainA_length <- 552

#Pick output file names
output_file <- "Links-out.txt"

pymol_output_file <- "PyMol-Swp1b-EnP1-XL.csv"

#Pick which protein is chain A and which is chain B
chain_A <- "EHEL_101700"
chain_B <- "EHEL_010690"


#filter for your proteins
df <- df[
  (df$Accession.A == chain_A & df$Accession.B == chain_B) |
    (df$Accession.A == chain_B & df$Accession.B == chain_A),
]

#Change accession number to chain letter
df$Accession.A[df$Accession.A == chain_A] <- "A"
df$Accession.B[df$Accession.B == chain_A] <- "A"
df$Accession.A[df$Accession.A == chain_B] <- "B"
df$Accession.B[df$Accession.B == chain_B] <- "B"

#Organize with chain A first
fixed_chain <- "A"
df <- df %>%
  mutate(
    # Check if the fixed chain is in column B
    swap = Accession.B == fixed_chain,
    
    # If swap = TRUE, flip A and B columns
    chain1 = ifelse(swap, Accession.B, Accession.A),
    res1   = ifelse(swap, In.protein.B, In.protein.A),
    chain2 = ifelse(swap, Accession.A, Accession.B),
    res2   = ifelse(swap, In.protein.A, In.protein.B)
  )

#If the residue is in chain B the residue will have chainA length added
df$res1[df$chain1 == "B"] <- df$res1[df$chain1 == "B"] + chainA_length
df$res2[df$chain2 == "B"] <- df$res2[df$chain2 == "B"] + chainA_length

#For chain B intralinks the length of chainA will be added
df_B$V1 <- as.numeric(df_B$V1) + chainA_length
df_B$V2 <- as.numeric(df_B$V2) + chainA_length


# Generate the formatted strings for crosslinks
output_lines <- apply(df, 1, function(row) {
  sprintf(
    "rsr.add(forms.upper_bound(group=physical.xy_distance,\n                               feature=features.Distance(at['CA:%s:%s'],\n                                                         at['CA:%s:%s']),\n                                  mean=%s, stdev=%s))",
    row["res1"], row["chain1"],  # first CA atom: V2 (resid), V1 (chain)
    row["res2"], row["chain2"],  # second CA atom: V5 (resid), V4 (chain)
    distance_cutoff,
    std_dev
  )
})

# Generate the formatted strings for intralinks in chain A
output_A <- apply(df_A, 1, function(row) {
  sprintf(
    "rsr.add(forms.upper_bound(group=physical.xy_distance,\n                               feature=features.Distance(at['CA:%s:A'],\n                                                         at['CA:%s:A']),\n                                  mean=%s, stdev=%s))",
    row[1], row[2],
    distance_cutoff, std_dev
  )
})

# Generate the formatted strings for intralinks in chain B
output_B <- apply(df_B, 1, function(row) {
  sprintf(
    "rsr.add(forms.upper_bound(group=physical.xy_distance,\n                               feature=features.Distance(at['CA:%s:B'],\n                                                         at['CA:%s:B']),\n                                  mean=%s, stdev=%s))",
    row[1], row[2],
    distance_cutoff, std_dev
  )
})

# Save crosslinks to a file
writeLines(output_lines, output_file)

# Append chain A intralinks
cat(paste0(output_A, "\n"), file = output_file, append = TRUE)

# Append chain B intralinks
cat(paste0(output_B, "\n"), file = output_file, append = TRUE)

### Make the file for compareMultimerDistances.py

#Prepare first dataframe
df_comb <- data.frame(
  chain1 = df$V1,
  chain2 = df$V4,
  residue1 = df$V2,
  residue2 = df$V5
)

#Prepare second dataframe
dfA_comb <- data.frame(
  chain1 = rep("A", nrow(df_A)),
  chain2 = rep("A", nrow(df_A)),
  residue1 = df_A$V1,
  residue2 = df_A$V2
)

#Prepare third dataframe
dfB_comb <- data.frame(
  chain1 = rep("B", nrow(df_B)),
  chain2 = rep("B", nrow(df_B)),
  residue1 = df_B$V1,
  residue2 = df_B$V2
)

df_all <- rbind(df_comb, dfA_comb, dfB_comb)

write.csv(df_all, pymol_output_file, row.names = FALSE)
```
